# Reconstitution of β-adrenergic regulation of Ca_V_1.2: Rad-dependent and Rad-independent protein kinase A mechanisms

**DOI:** 10.1101/2020.11.30.403964

**Authors:** Moshe Katz, Suraj Subramaniam, Orna Chomsky-Hecht, Vladimir Tsemakhovich, Anouar Belkacemi, Veit Flockerzi, Enno Klussmann, Joel A. Hirsch, Sharon Weiss, Nathan Dascal

## Abstract

**Introduction:** Cardiac L-type voltage-gated Ca_V_1.2 channels are crucial in physiological regulation of cardiac excitation-contraction coupling. Adrenergic modulation of Ca_V_1.2 starts with activation of β-adrenergic receptors (AR) and culminates in protein kinase A (PKA) - induced increase of calcium influx through Ca_V_1.2 channels. To date, this cascade has never been fully reconstituted in heterologous systems; even partial reconstitution proved challenging and controversial. A recent study identified Rad, a calcium channel inhibitory protein, as an essential component of the adrenergic signaling cascade. We corroborated this finding, further characterized, and fully reconstituted, the complete β-AR Ca_V_1.2 modulation cascade in a heterologous expression system.

**Objective:** Our primary goal was to heterologously reconstitute the complete β-adrenergic cascade, and to investigate the role of Rad and additional molecular determinants in adrenergic regulation of cardiac Ca_V_1.2.

**Methods and Results:** We utilized the *Xenopus* oocyte heterologous expression system. We expressed Ca_V_1.2 channel subunits, without or with Rad and β1-AR or β2-AR. To activate PKA, we injected cyclic AMP (cAMP) into the oocytes, or extracellularly applied isoproterenol (Iso) to stimulate β-AR. Whole-cell Ba^2+^ currents served as readout. We find and distinguish between two distinct pathways of PKA modulation of Ca_V_1.2: Rad-dependent (~80% of total) and Rad-independent. We separate the two mechanisms by showing distinct requirements for the cytosolic N- and distal C- termini of α_1C_ and for the Ca_V_β subunit. Finally, for the first time, we reconstitute the complete pathway using agonist activation of either β1-AR or β2-AR. The reconstituted system reproduces the known features of β-AR regulation in cardiomyocytes, such as a >2-fold increase in Ca_V_1.2 current, a hyperpolarizing shift in activation curve, and a high constitutive activity of β2-AR.

**Conclusions:** The adrenergic modulation of Ca_V_1.2 is composed of two distinct pathways, Rad-independent and Rad-dependent. The latter contributes most of the β-AR-induced enhancement of Ca_V_1.2 activity, crucially depends on Ca_V_β subunit, and is differently regulated by β1-AR and β2-AR. The reconstitution of the full β-AR cascade provides the means to address central unresolved issues related to roles of auxiliary proteins in the cascade, Ca_V_1.2 isoforms, and will help to develop therapies for catecholamine-induced cardiac pathologies.

## Introduction

Cardiac excitation-contraction coupling crucially depends on the L-type voltage dependent Ca^2+^ channel, Ca_V_1.2. Influx of extracellular Ca^2+^ via Ca_V_1.2 triggers Ca^2+^ release from the sarcoplasmic reticulum, via the Ca^2+^ release channel (ryanodine receptor 2)^1^. Activation of the sympathetic nervous system increases heart rate, relaxation rate and contraction force. The latter is due largely to increased Ca^2+^ influx via Ca_V_1.2^2^. Pathological prolonged sympathetic activation progressively impairs cardiac function, causing heart failure, partly due to misregulation of Ca_V_1.2^3^.

Cardiac Ca_V_1.2 is a heterotrimer comprising the pore-forming subunit α_1C_ (~240 kDa), the intracellular Ca_V_β_2_ (~68 kDa) and the extracellularly located α2δ (~170 kDa)^4, 5^. The N and C termini (NT, CT respectively) of α_1C_ are cytosolic and vary among Ca_V_1.2 isoforms (Fig. 1A). It is believed that most of the cardiac α_1C_ protein is post-translationally cleaved at the CT, around amino acid (a.a.) 1800, to produce the truncated ~210 kDa α_1C_ protein and the ~35 kDa cleaved distal CT (dCT); however, the full-length protein is also present^6–9^.

**Figure 1.**
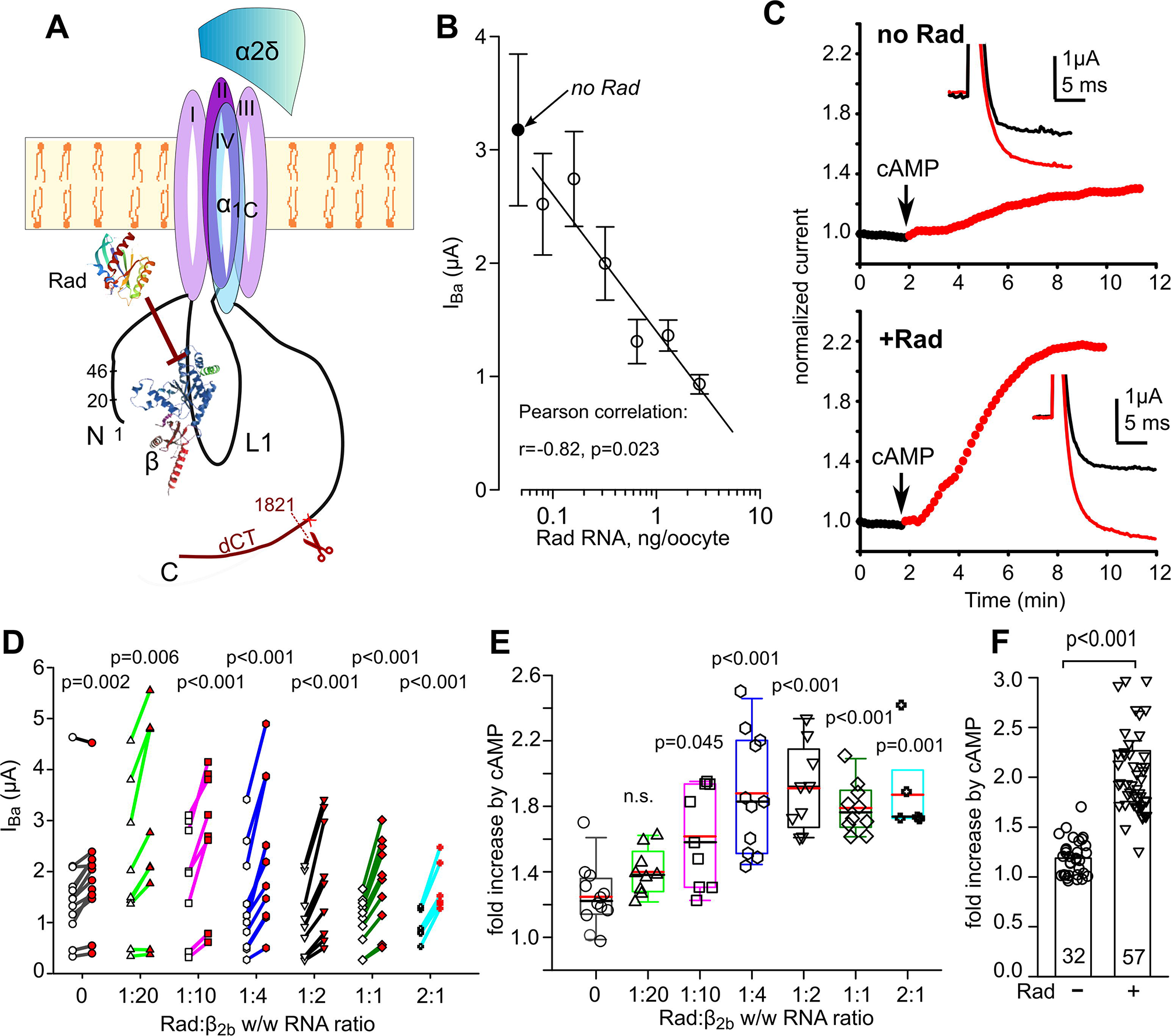
cAMP regulation of Ca_V_1.2 is greatly enhanced by co-expression of Rad. **A**, The cardiac Ca_V_1.2 and Rad. The α_1C_ and α2δ subunits are shown schematically, along with structures of β_2b_ (PDB:1T3S)^30^ and Rad (PDB: 3Q72)^33^. In α_1c_, the four homologous transmembrane domains are numbered I to IV. The constructs used here were based on rabbit or mouse long-NT isoforms, containing the 46 a.a.-long initial segment encoded by exon 1a^34^ and either full-length CT, or, in α_1C_Δ1821, the α_1C_ was truncated at a.a. 1821, close to the naturally occurring truncation site (red cross mark) in cardiac α_1c_, ~a.a. 1800^7^. β_2b_ binds to the cytosolic loop connecting domains I and II. When recruited to the plasma membrane, Rad interferes with the interaction between β and α_1C_ subunits. **B-F**, examples and summaries for Ca_V_1,2-α_1C_Δ1821. **B**, Rad reduces the Ba^2+^ current of Ca_V_1.2-α_1C_Δ1821 in a dose-dependent manner. The channel was expressed in full subunit composition: α_1C_Δ1821, β_2b_ and α2δ (1.5 ng RNA of each subunit). I_Ba_ decreased with increasing doses of Rad RNA (Pearson correlation, r=− 0.82, p=0.023). Each point represents mean±SEM from 7 to 10 oocytes (N=1 experiment). The linear regression line was drawn for non-zero doses of Rad. C, Rad enhances the cAMP-induced increase in I_Ba_. Figures are diary plots showing the time course of change in I_Ba_ (normalized to initial I_Ba_), before and after the intracellular injection of cAMP in a representative cell. Insets show current records at +20 mV before (black trace) and 10 min after cAMP injection (red trace), without Rad in the upper panel and with Rad in the lower panel. **D**, “before-after” plots of cAMP-induced changes in I_Ba_ in individual cells injected with increasing ratio of Rad:β_2b_ expression. Empty symbols – before cAMP; red-filled – after cAMP. RNA ratios (by weight, w/w) of Rad and β_2b_ were varied in 1:20 to 2:1 range. N=3 experiments; statistics: paired t-test. **E**, cAMP-induced increase in I_Ba_ depends on relative expression levels of Rad and β_2b_. Each symbol represents fold increase in I_Ba_ induced by cAMP injection in an individual cell, at different Rad:β_2b_ RNA ratios. Here and in the following figures, box plots show 25-75 percentiles, whiskers show the 5/95 percentiles, the black and the red horizontal lines within the boxes are the median and mean, respectively. At all Rad:β_2b_ RNA ratios except 1:20, the cAMP-induced increase in I_Ba_ was significantly greater than without Rad (Kruskal-Wallis One-Way ANOVA on ranks). **F**, summary of the effect of cAMP in 10 experiments without Rad and with Rad at 1:2 and 1:1 Rad:β_2b_ RNA ratios (pooled). Number of cells is shown within the bars. Statistics: Mann-Whitney test.

The sympathetic nervous system activates cardiac β-adrenergic receptors (β-AR), primarily β1-AR (which mediates most of the βAR-enhancement of contraction and Ca_V_1.2 activity) and β2-AR^10^. β1-AR is globally distributed in cardiomyocytes, couples to G_s_ and elevates cAMP levels within the whole cardiomyocyte^10^. In contrast, β2-AR shows spatially restricted localization to the T-tubules (which changes in heart failure)^11^ and couples to G_s_ and G_i/o_ ^12^, producing localized cAMP increases^3^.

The cascade of adrenergic modulation of Ca_V_1.2 comprises agonist binding to β-ARs, activation of G_s_, adenylyl cyclase, elevated intracellular cAMP levels, and activation of protein kinase A (PKA). The PKA holoenzyme consists of two regulatory subunits (PKA-RS) bound to two catalytic subunits (PKA-CS). cAMP binds to PKA-RS and causes dissociation of PKA-CS and PKA-CS. The latter enhances Ca_V_1.2 activity. However, the final step, how PKA-CS enhances Ca_V_1.2 activity, remained enigmatic. A long standing paradigm was a direct phosphorylation by PKA-CS of α_1C_ and/or Ca_V_β subunits^2^. However, numerous studies critically challenged this theory. In particular, mutated Ca_V_1.2 channels in genetically engineered mice lacking putative PKA phosphorylation sites on α_1C_ and/or β_2b_, were still upregulated by PKA^7, 13–16^ (reviewed in^4, 17^).

One significant obstacle in deciphering the mechanism of PKA regulation of Ca_V_1.2 was a recurrent failure to reconstitute the regulation in heterologous systems, which proved challenging and controversial^18^. Previous studies in heterologous cellular models, including *Xenopus* oocytes, demonstrated that cAMP failed to up-regulate Ca_V_1.2 containing the full-length α_1C_, Ca_V_1.2-α_1C_^19–21^. However, robust β-AR – induced upregulation of Ca^2+^ currents was observed in oocytes injected with total heart RNA^22^, suggesting the necessity of an auxiliary protein, the “missing link”^19, 20^. Interestingly, partial regulation was observed with dCT-truncated α_1C_^23, 24^, which is considered the predominant form of α_1C_ in the heart^2^. We have previously reported a modest (30-40%) upregulation of Ca_V_1.2, containing a dCT-truncated α_1C_, by intracellular injection of cAMP or PKA-CS in *Xenopus* oocytes ^24^. Additionally, this regulation required the presence of the initial segment of the long-NT of α_1C_, but did not involve Ca_V_β subunit. We proposed that this mechanism may account for part of the adrenergic regulation of Ca_V_1.2 in the heart^24^. Normally adrenergic stimulation in cardiomyocytes increases the Ca^2+^ current two to three fold, thus a major part of the regulation has remained unexplained.

Recently, Liu et al. identified Rad as the “missing link” in PKA regulation of Ca_V_1.2^15^. Rad is a member of the Ras-related GTPase subfamily (RGK) that inhibits high voltage activated calcium channels Ca_V_1 and Ca_V_2^25^. Rad tonically inhibits Ca_V_1.2, largely via an interaction with Ca_V_β^26, 27^. Ablation of Rad in murine heart was shown to increase basal Ca_V_1.2 activity but suppressed β-AR regulation^28^. Liu et al. demonstrated that PKA phosphorylation of Rad relieves this tonic inhibition to increase Ca_V_1.2 activity^15^. They reconstituted a major part of the Ca_V_1.2 regulation cascade, initiated by forskolin-activated adenylyl cyclase and ultimately attaining a ~2-fold increase in Ca^2+^ current, in mammalian cells expressing α_1C_, α2δ, β_2b_ and Rad^15^. The relation between this Rad-dependent regulation and the regulation reported in our previous study^24^ is not clear. Furthermore, the complete pathway of adrenergic modulation of Ca_V_1.2, starting with β-AR activation, has not yet been heterologously reconstituted.

Here we utilized the *Xenopus* oocyte heterologous expression system and, for the first time, reconstituted the entire pathway. We find and distinguish between two distinct pathways of PKA modulation of Ca_V_1.2: a Rad-dependent and a Rad-independent pathway. We characterize the involvement of the N- and C- termini of α_1C_ and of the β_2b_ subunit in this crucial physiological regulation, and compare and contrast the Ca_V_1.2 regulation by β1-AR or β2-AR. Our findings reveal novel aspects of the roles of α_1C_ (particularly the N- and C-termini), β_2b_ and Rad in the adrenergic modulation of cardiac Ca_V_1.2 channels. Reproducing the complete β-AR cascade in a heterologous expression system will promote the identification and characterization of intracellular proteins that regulate the cascade, eventually assisting efforts to develop therapies to treat heart failure and other catecholamine-induced cardiac pathologies.

## Methods

### Experimental animals and ethical approval

Oocytes were harvested from adult female *Xenopus laevis* frogs, as described^24^. Ethical approval was granted by Tel Aviv University Institutional Animal Care and Use Committee (permits 01-16-104 and 01-20-083).

### DNA constructs and RNA

In this work we expressed Ca_V_1.2 in *Xenopus* oocytes, usually in full subunit composition α_1C_+α2δ+β_2b_, by injection of equal amounts, by weight, of RNAs of each subunit (except experiments of Fig. 2 that addressed the role of the Ca_V_β subunit). We used the long-NT isoform of rabbit α_1C_ (except Fig. S1 where mouse α_1C_ was used) and various mutants, as detailed in the figures. RNAs of β1-AR, β2-AR, human Rad, or the α_1C_ dCT (as a separate protein) were expressed according to the design of the experiment. The Ca_V_β subunit used here was rabbit β_2b_ (originally termed β_2a_^29^). We tested the full-length β_2b_ and β_2b_-core, composed of a.a. 26-422 of Ca_V_β with deletion of the linker sequence (a.a. 138-202) as previously described^30^. The β_2b_-core_3DA_ mutant^31^ was prepared based on the β_2b_-core construct with three point mutations (D244A/D320A/D322A) (see Fig. 2A). RNAs of all constructs were prepared and injected into oocytes as described^24^.

**Figure 2.**
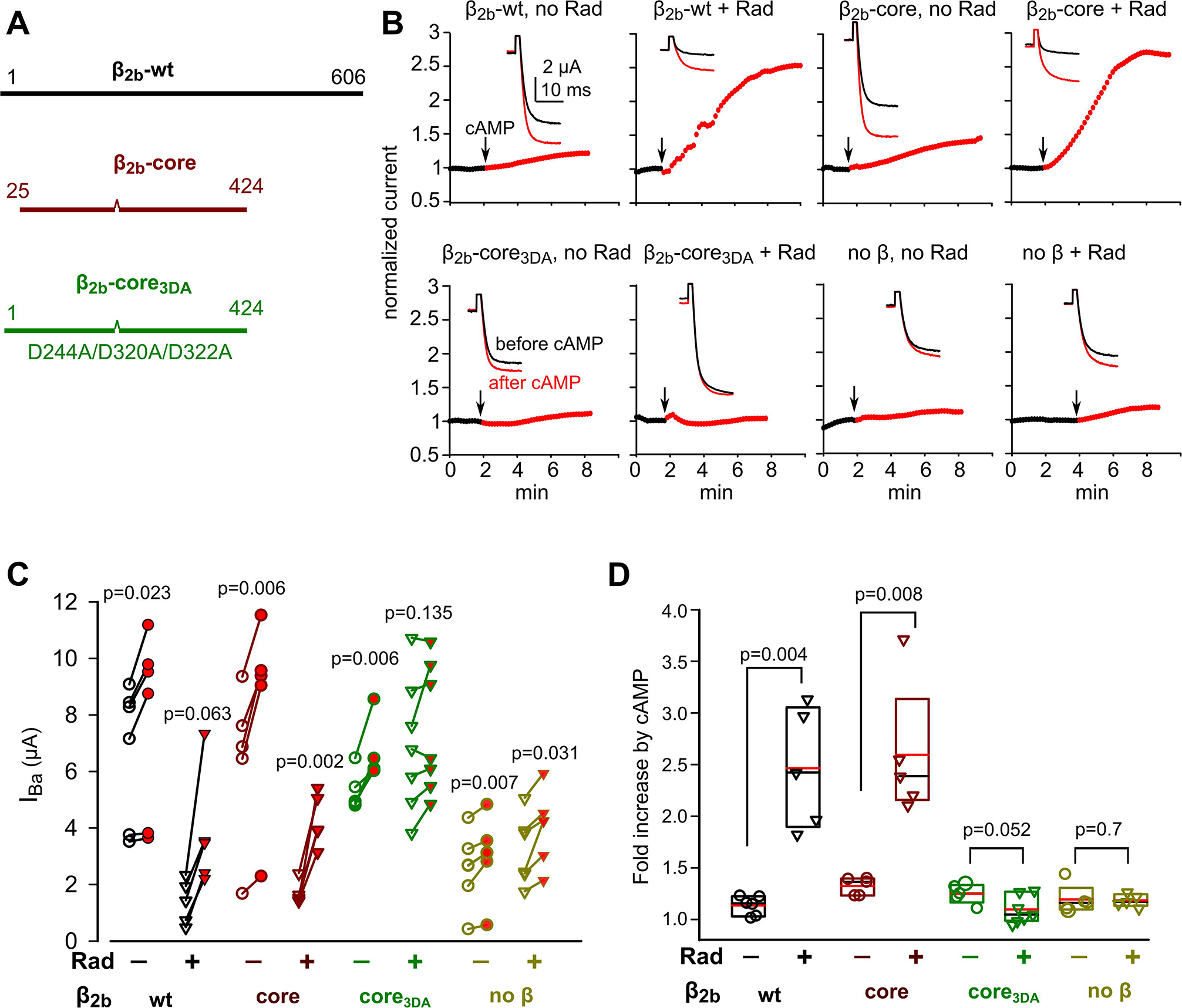
Separation of Rad-dependent and Rad-independent PKA regulation of α_1C_: the role of Ca_V_β. **A**, Schematic representation of Ca_V_β subunit variants used in this study. The wild-type (β_2b_-wt) protein is 632 a.a. long. β-core is truncated at a.a. 424; the linker amino acids 138-202 were removed^30^. The β-core_3DA_ is the β-core with three aspartate-to-alanine mutations, D244A/D320A/D322A, that does not bind Rad.. **B-D**, the presence of the β subunit and its ability to bind Rad are crucial for Rad-dependent, but not for Rad-independent, cAMP regulation of Ca_V_1.2. Rad:β_2b_ RNA ratio was 1:1. N=1 experiment. **B**, Diary plots of cAMP-induced changes in I_Ba_ following injection of cAMP in representative cells expressing α_1C_Δ1821, α2δ, and the indicated variant of β_2b_ (or no β at all), with or without Rad. Insets show current records at +20 mV before (black trace) and 10 min after cAMP injection (red trace). **C**, “Before-after” plots of cAMP-induced changes in I_Ba_ in individual cells injected with various Ca_V_β subunits co-expressed with and without Rad. N=1 experiment; statistics: paired t-test. **D**, Fold change in I_Ba_ after cAMP injection. Data show the fold increase with Rad co-expression (inverted triangles) and without Rad (circles). Mann-Whitney Rank Sum test was used to compare groups with and without Rad (except the groups with β-core_3DA_ where normality was satisfied, and t-test was used).

### Electrophysiology

Recordings were performed on defolliculated oocytes 3-4 days post RNA injection. Whole-cell Ba^2+^ currents via Ca_V_1.2 (I_Ba_) were recorded using two-electrode voltage clamp, routinely in a 40 mM Ba^2+^ solution (in mM: 40 Ba(OH)_2_,50 NaOH, 2 KOH, and 5 Hepes, titrated to pH 7.5 with methanesulfonic acid). The membrane potential was −80 mV and I_Ba_ was elicited by steps to +20 mV for 20 ms every 10 s. To obtain the current-voltage (I–V) relationship, currents were elicited by 20 ms pulses starting at −70 mV up to +80 mV in 10 mV increments every 10 s. Currents measured in the presence of 200 μM Cd^2+^ were subtracted from total I_Ba_ (Fig. 5A-C) to yield the net I_Ba_. I-V curves were fitted using the Boltzmann equation^32^. The parameters obtained for G_max_ and V_rev_ were then used to calculate fractional conductance at each V_m_ and to construct the conductance-voltage (G-V) curve. CFTR currents were measured at −80 mV in ND96 solution (in mM: 96 NaCl, 2 KCl,1 MgCl_2_, 1 CaCl_2_, 5 Hepes, pH 7.6).

cAMP, PKA or Iso were applied after verifying that I_Ba_ is stable for at least 2 min. cAMP or PKA-CS were introduced into oocytes by pressure microinjection to final concentrations within the oocyte of ~100 μM (cAMP) or ~25 ng/oocyte (PKA-CS). Isoprenaline hydrochloride (isoproterenol, Iso) was perfused into the experimental chamber.

### Statistical analysis

Results are presented as median and interquartile range (IQR) [Q1-Q3], or as mean ± standard error for normally distributed continuous variables. For comparisons between groups, we used Student’s t-test or Mann-Whitney test, as appropriate. The fold change in current caused by a treatment in a single oocyte was calculated as (I_Ba_ after treatment)/(I_Ba_ before treatment). The I_Ba_ amplitudes before and after treatment in a single cell were compared using paired t-test or Wilcoxon test, as appropriate. For multiple group comparisons, we performed a One-Way ANOVA or Kruskal Wallis ANOVA on ranks, as appropriate. A Bonferroni post hoc test was performed for normally distributed data (Shapiro-Wilk test) and Dunnett’s post hoc test otherwise. Statistical analysis was performed with SigmaPlot 13 (Systat Software Inc., San Jose, CA, USA).

## Results

### Rad plays a significant role in PKA regulation of Ca_V_1.2

PKA regulation of heterologously expressed Ca_V_1.2 was previously observed only with dCT-truncated, but not full-length α_1C_ ^23, 24^. In *Xenopus* oocytes, Ba^2+^ currents (I_Ba_) via Ca_V_1.2 containing dCT-truncated α_1C_ (Ca_V_1.2-α_1C_Δ1821) were increased by 20-40% following intracellular injection of cAMP or PKA-CS^24^. Therefore, we started the study of Rad’s role in regulation of Ca_v_1.2 using α_1C_Δ1821.

We expressed Ca_V_1.2 in full subunit composition, α_1C_Δ1821, β_2b_ and α2δ, unless indicated otherwise, at a 1:1:1 RNA ratio (by weight), and tested the effect of increasing Rad concentrations by injecting varying amounts of Rad RNA. I_Ba_ was measured 3-5 days after RNA injection, by voltage steps from −80 to +20 mV (see Fig. 1C, inserts, for examples of I_Ba_ recordings). As expected^26, 31^, basal I_Ba_ was reduced by expression of Rad, showing an inverse correlation with Rad RNA dose (Fig. 1B). Without Rad coexpression, I_Ba_ was increased by 19%±3 (n=32, p=0.002) following cAMP injection (Fig. 1C-E). Coexpression of Rad dramatically augmented the effect of cAMP (Fig. 1C, E). This effect of Rad became statistically significant at Rad:β_2b_ RNA ratios above 1:10, with a maximum increase of 127±10% at 1:2 ratio (Fig. 1F).

### *The role of Ca*_*V*_β *in Rad dependent regulation*

Rad inhibits Ca_V_1.2 through Ca_V_β-dependent and Ca_V_β-independent mechanisms^26, 31^. To test the role of Ca_V_β in PKA regulation of Ca_V_1.2, we compared the change in I_Ba_ following injection of cAMP into oocytes co-expressing α_1C_Δ1821, α2δ, and β_2b_-wt (wild-type), β_2b_-core^30^, β_2b_-core_3DA_ or no β_2b_, with or without Rad (Fig. 2A). β_2b_-core_3DA_ was designed on the basis of β_2b_-core and contained the triple mutation D244A/D320A/D322A which abolishes Ca_V_β-Rad association^31, 35^.

In oocytes that did not express Rad, cAMP induced the typical, mild but statistically significant, 20-30% increase in I_Ba_ (Fig. 2B-D). This regulation is termed hereafter “Rad-independent”. The Rad-independent cAMP effect was similar in oocytes that did not express any Ca_V_β or expressed β_2b_, β_2b_-core, or the β_2b_-core_3DA_ mutant (Fig. 2B-D). Co-expression of Rad resulted in a much larger, ~2.5-fold increase in peak currents, but only when β_2b_-wt or β_2b_-core were present (Fig. 2B, 2D). We refer to this regulation as “Rad-dependent”.

Importantly, in Rad-expressing oocytes that did not express Ca_V_β, or expressed the β_2b_-core_3DA_ mutant, cAMP caused only a mild increase in I_Ba_ that resembled the Rad-independent cAMP effect (Fig. 2B-D). Notably, basal I_Ba_ was significantly increased by coexpression of β_2b_-core_3DA_ (e.g. Fig. 2C) indicating robust protein expression and channel regulation. Evidently, abrogation of β_2b_-Rad interaction eliminates the Rad-dependent PKA regulation of Ca_V_1.2. Thus, Rad-dependent PKA regulation of Ca_V_1.2 is Ca_V_β-dependent. β_2b_-core is sufficient to mediate the regulation. In contrast, the Rad-independent mechanism does not require Ca_V_β (as shown before^24^) and its contribution to cAMP-induced current increase is much smaller than that of the Rad-dependent one.

### The N-terminus of α_1C_ is important for Rad-independent but not Rad-dependent regulation

We then set out to characterize the determinants of Rad-independent regulation. The predominant cardiac isoform is the long-NT α_1C_, where the first 46 a.a., out of ~154, are encoded by the alternative exon 1a^34^. The initial segment (first 20 a.a.) of this α_1C_ isoform acts as an inhibitory module, tonically reducing the activity of Ca_V_1.2 by reducing the channel’s open probability^32^. Within the NT initial segment, a.a. 6-20 show partial homology with the first 16 a.a. of the short NT isoform (these 16 a.a. are encoded by the alternative exon 1), found in smooth muscle and brain^4^ (Fig. S1A). Specifically, a.a. T_10_, Y_13_ and P_15_ (TYP motif) are highly conserved and crucial for the inhibitory function of the NT module^32^. Importantly, the Rad-independent PKA regulation of Ca_V_1.2-α_1C_Δ1821 is greatly reduced or abolished by the removal of the first 5 or 20 a.a. of the long NT^24^. We have further addressed the role of the NT initial segment by expressing Ca_V_1.2 with mouse α_1C_Δ1821 containing alanine substitutions of a.a. 2-5 (α_1C_NT-4AΔ1821) or the TYP motif (α_1C_NT-TYPΔ1821) (Fig. S1, B-D). PKA-CS injection into oocytes expressing α_1C_Δ1821 with intact NT increased I_Ba_ by ~70% (p=0.004). In contrast, oocytes expressing α_1C_NT-4AΔ1821 or α_1C_NT-TYPΔ1821 did not respond to PKA-CS. Thus, as shown previously^24^, the initial segment of the long-NT-α_1C_, including the first 5 a.a. and the TYP motif, is essential for the Rad-independent PKA regulation of cardiac Ca_V_1.2.

To further address the role of the NT of α_1C_Δ1821 in Rad-independent vs. Rad-dependent PKA regulation of Ca_V_1.2, we used NT-truncated-α_1C_Δ1821 with deletion of the first 20 a.a. (α_1C_Δ20Δ1821) (see Fig. 1A). We verified that the Rad-independent increase in I_Ba_ of α_1C_Δ1821 (34.4±7.5%; n=10) was greatly diminished in α_1C_Δ20Δ1821 (5±2%, n=16; p<.001 compared to α_1C_Δ1821) (Fig. 3A, B). In contrast, co-expression of Rad resulted in similar ~1.7-2 fold increases in I_Ba_ following cAMP injection when Ca_V_1.2 contained either α_1C_Δ1821 or α_1C_Δ20Δ1821 (p=0.154). In both constructs, the fold increase in I_Ba_ was statistically significant in the presence of Rad, and significantly greater than without Rad (Fig. 3). Thus, Rad-dependent regulation does not require the presence of the NT inhibitory module.

**Figure 3.**
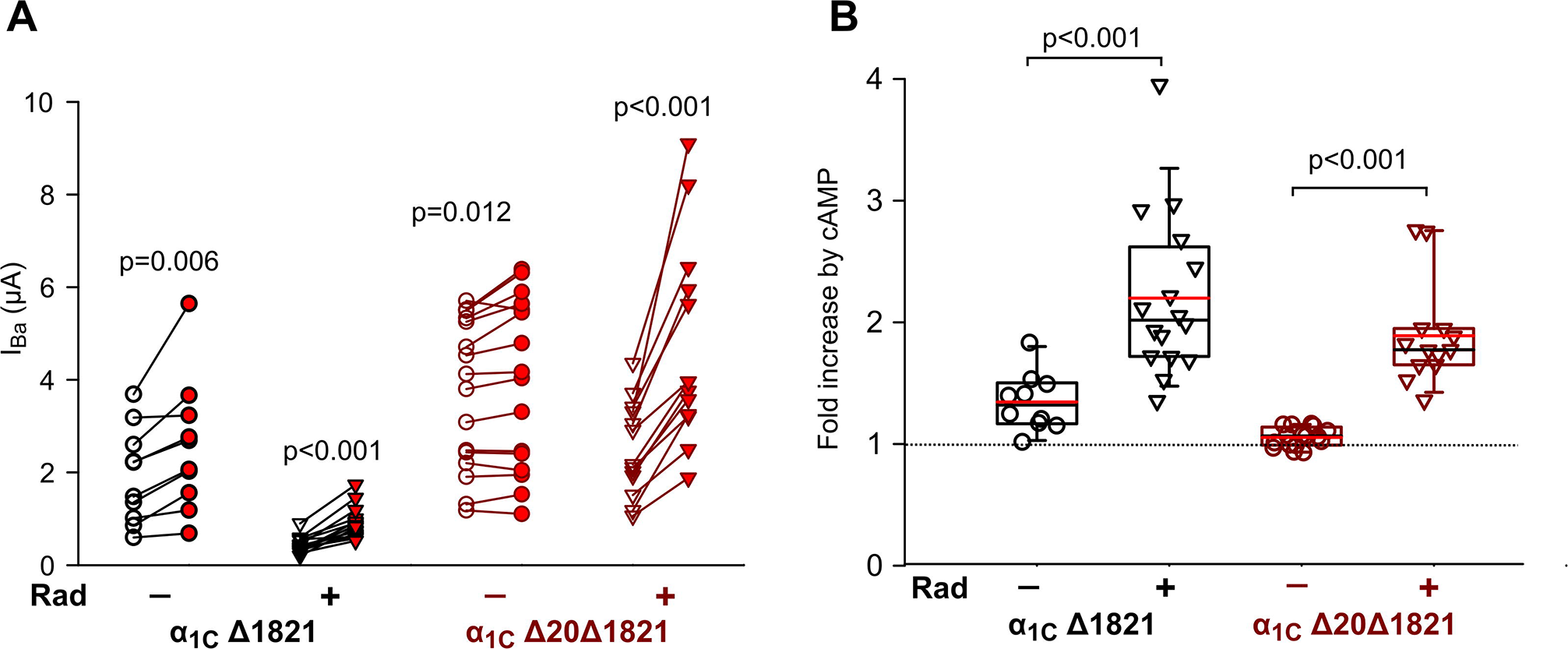
Separation of Rad-dependent and Rad-independent PKA regulation of α_1C_: the role of N-terminal initial segment of α_1C_. **A**, “before-after” plots of cAMP-induced changes in I_Ba_ in individual cells expressing Ca_V_1.2 containing α_1C_Δ1821 (black) and α_1C_ Δ20Δ1821 (red) lacking the first 20 a.a. of the N-terminus. α2δ and β_2b_ were co-expressed in all cases, without or with Rad (circles and inverted triangles, respectively). N=3 experiments; statistics: paired t-test. **B**, Fold change in I_Ba_ after cAMP injection (summary of experiments shown in A). The black and the red horizontal lines within the boxes are the median and mean, respectively. Pairwise comparison with and without Rad was done using the Mann-Whitney Rank Sum test.

### The role of distal C-terminus of α_1C_ in PKA regulation

It has been proposed that the cleaved dCT is a potent autoinhibitory domain that reassociates with the truncated α_1C_, forming a tight molecular complex^8^ that is essential for PKA regulation of Ca_V_1.2^2, 23^. In addition, the cleaved dCT has been reported to traffic to the nucleus where it serves as a transcription regulator^36, 37^. However, in *Xenopus* oocytes, the presence of dCT as a separate protein was not required for Rad-independent regulation of Ca_V_1.2-Δ1821^24^. Recently, forskolin-induced upregulation of Ca_V_1.2 was demonstrated in HEK293T cells co-expressing Rad and full-length Ca_V_1.2 channels^15^. All in all, the role of dCT and its truncation in Rad-dependent regulation remains incompletely understood, and it is unknown whether this PKA regulation equally affects full-length and truncated forms of α_1C_.

To examine the role of dCT in Rad-dependent PKA regulation of Ca_V_1.2, we compared the effect of cAMP on either full length (wt) α_1C_ or α_1C_Δ1821 channels, in the absence and presence of Rad. We also examined the effect of dCT when coexpressed as a separate protein. I_Ba_ is greatly increased by the truncation of dCT^38^. Therefore, to maintain similar macroscopic currents, we injected different amounts of the channel’s subunit RNAs: 1-1.5ng RNA for Ca_V_1.2 containing α_1C_Δ1821, and 1.8-5 ng RNA for Ca_V_1.2 containing wt-α_1C_. The dCT:α_1C_Δ1821 RNA ratio was 5:1 and Rad:β_2b_ RNA ratios were in the range of 1:3 to 1:1 (which yielded similar increase in I_Ba_ when cAMP is injected; see Fig. 1E). As with α_1C_Δ1821, co-expression of Rad reduced basal currents of Ca_V_1.2 containing wt-α_1C_, with median I_Ba_ of 3.05 μA [IQR 2.46-4.73] without Rad and 0.62 μA [IQR 0.47-0.73] with Rad (p<.001; Fig. 4B).

**Figure 4.**
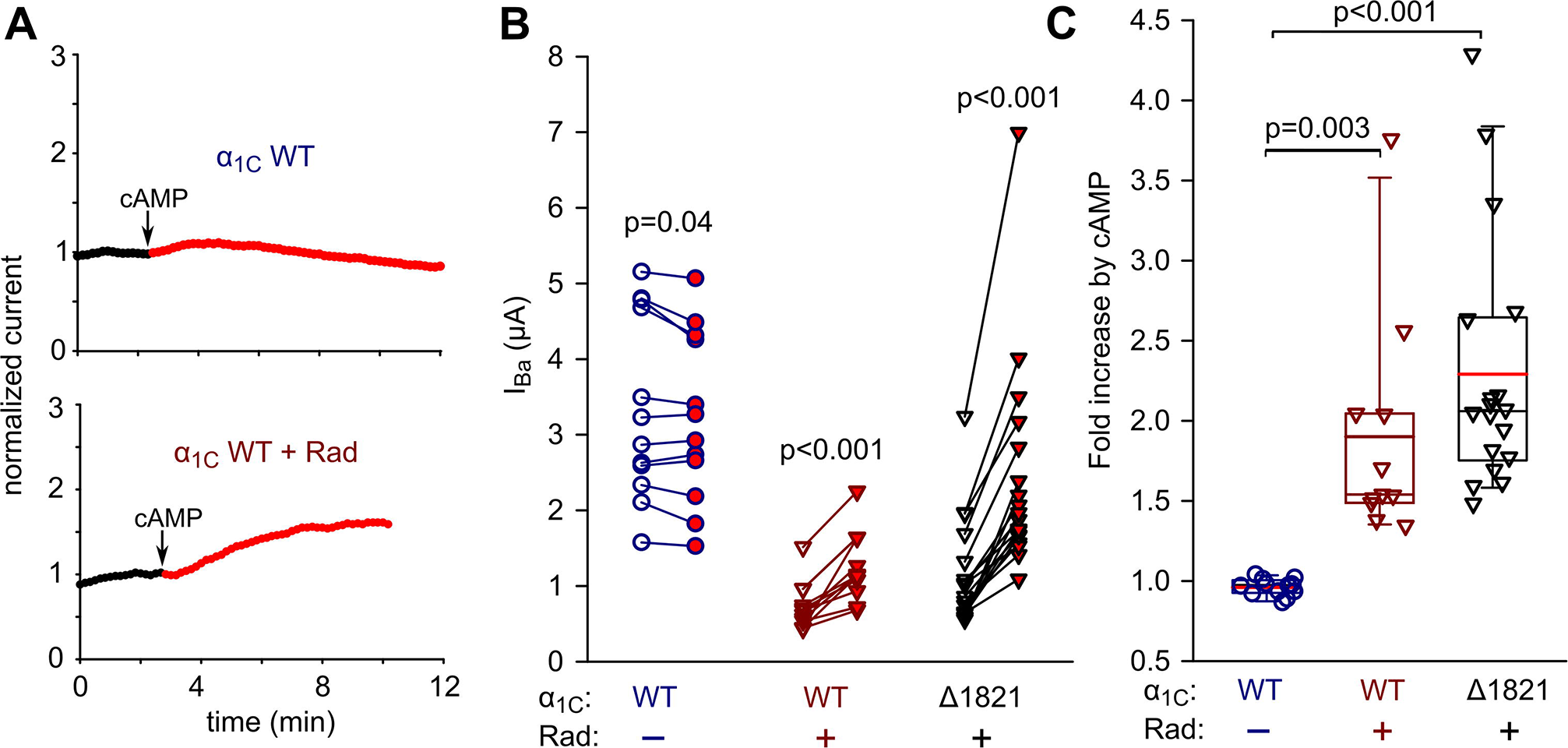
Separation of Rad-dependent and Rad-independent PKA regulation of α_1C_: the role of the distal C-terminus of α_1C_. **A**, Diary plots of cAMP-induced changes in I_Ba_ in representative cells expressing the full length α_1C_ (α_1C_ WT), α2δ and β_2b_, without Rad (upper panel) or with Rad (lower panel). **B**, “before-after” plots of cAMP-induced changes in I_Ba_ in individual cells expressing wt-α_1c_ with or without Rad co-expression, or α_1C_Δ1821 and Rad. α2δ and β_2b_ were present in all cases. Rad:β_2b_ RNA ratio was 1:2 or 1:1. Statistics: paired t-test. N=3 experiments. n=12, 11 and 18 oocytes, from left to right. **C**, Fold change in I_Ba_ at +20 mV after cAMP injection (summary of data shown in A, B). The black and the red horizontal lines within the boxes are the median and mean, respectively. Statistics: Kruskal-Wallis One-Way ANOVA on ranks.

Injecting cAMP into cells expressing wt-α_1C_ did not increase I_Ba_ (actually, a slight reduction of 4±1.5%, n=12, was observed: Fig. 4A, upper panel; Fig. 4B, C). In contrast, when Rad was coexpressed with wt-α_1C_, cAMP injection resulted in statistically significant increase of 90±21% in I_Ba_ (n=11) (Fig. 4A, lower panel; Fig. 4B, C). Interestingly, in the same experiments, the Rad-dependent cAMP-induced increase in I_Ba_ appeared higher in oocytes co-expressing Rad with α_1C_Δ1821: 128±23%, n=18. Pairwise comparison of fold increase in I_Ba_ for Ca_V_1.2 with wt-α_1C_ vs. α_1C_Δ1821 showed a mildly significant difference, p=0.04 (Mann-Whitney test, median 2.06 [IQR 1.77-2.63] for α_1C_Δ1821 vs. median 1.54 [IQR 1.49-2.04] for wt-α_1C_). When dCT was co-expressed as a separate protein with α_1C_Δ1821, it did not affect regulation by cAMP even in the presence of Rad (p=0.48) (Fig. S2). Thus, unlike Rad-independent regulation, Rad-dependent PKA regulation does not require the cleavage of dCT. Furthermore, the clipped dCT does not appear to play a role in Rad-dependent PKA regulation of α_1C_Δ1821. However, there appears to be a quantitative difference in the overall cAMP regulation of full-length versus truncated Ca_V_1.2.

### Full reconstitution of the β1 adrenergic receptor regulation of Ca_V_1.2

Despite more than a 3-decade effort, it has not yet been possible to reconstitute the entire adrenergic regulation of cardiac Ca_V_1.2 in a heterologous model. Here we report the reconstitution of the full cascade, starting with activation of β1-AR. The initial experiments were conducted with Ca_V_1.2 containing α_1C_Δ1821, using a Rad:β_2b_ RNA ratio of 1:2. In the absence of Rad, isoproterenol (Iso; 50 μM), a non-selective β-AR agonist^39^, did not produce any significant increase in I_Ba_ (Fig. 5D, E; Fig. 6). However, co-expression of Rad resulted in a significant increase in I_Ba_ following Iso application. Fig. 5A-B shows traces of currents (Fig. 5A) and current-voltage relationship (Fig. 5B) in a representative oocyte. Fig. 5C shows a conductance-voltage curve drawn from 7 oocytes of the same day’s experiment. Iso not only increased currents amplitudes, but also caused a ~5 mV hyperpolarization shift in the V_1/2_ for activation, without changing the slope factor (Fig. 5B, lower panel).

**Figure 5.**
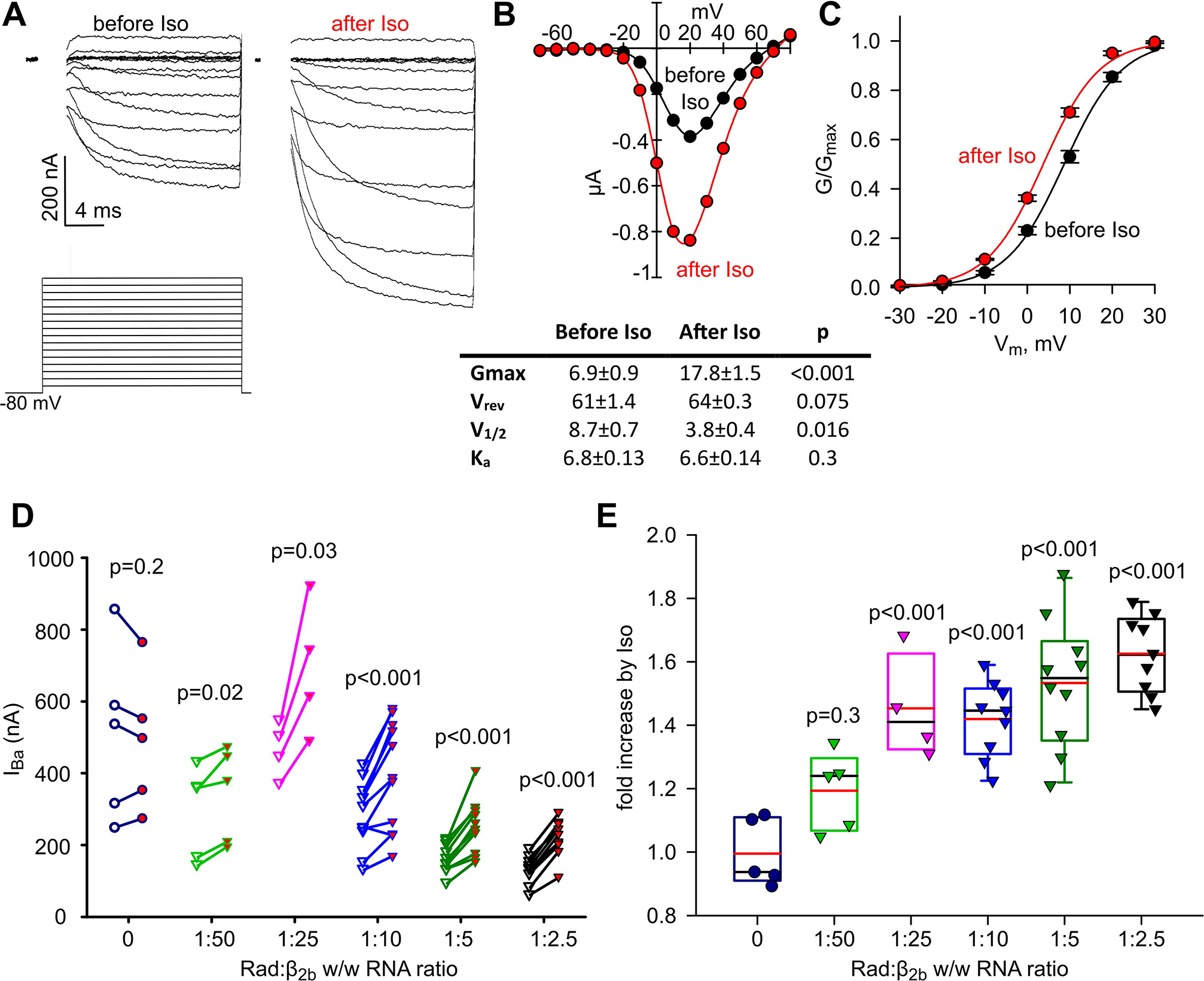
Full reconstitution of the β1-AR regulation of α_1C_. **A-C**, β1-AR regulation of Ca_V_1.2. Oocytes were injected with RNA of α_1C_Δ1821, α2δ, β_2b_, Rad and β1-AR. **A**, Ba^2+^ currents (upper panels) before (left) and after (right) perfusion of 50 μM isoproterenol (Iso) in a representative cell. The voltage protocol is illustrated in the lower panel; I_Ba_ was elicited by 20-ms voltage steps given every 10 s from a holding potential of −80 mV in 10 mV increments. The currents shown are net I_Ba_ derived by subtraction of the residual currents recorded with the same protocols after applying 200 μM Cd^2+^ (not shown). Since full capacity compensation in oocytes was not achievable, the currents during the first ~2 ms (the duration of capacity transient) were blanked out. **B**, Top, representative I-V curve before (black) and after (red) addition of Iso. Bottom, parameters of Boltzmann fit of I-V curves in 7 oocytes, before and after Iso. **C**, Conductance-voltage (G-V) curves of Ca_V_1.2-α_1C_Δ1821 co-expressed with Rad and β1-AR averaged from oocytes of a representative batch (n=7 oocytes, N=1 experiment) before and after Iso. The curves were drawn using the Boltzmann equation using average V_1/2_ and K_a_ obtained from the fits of I-V curves in individual oocytes (see Methods). Parameters used were: V_1/2_=8.7 mV, K_a_=6.8 mV before Iso; V_1/2_=3.8 mV, K_a_=6.6 mV after Iso. **D-E**, β1-AR regulation of Ca_V_1.2 containing the wt-α_1C_ in the presence of increasing doses of Rad. Oocytes were injected with RNAs of wt-α_1C_, α2δ, β_2b_, β1AR, and the indicated doses of Rad RNA. **D**, “before-after” plots of Iso-induced changes in I_Ba_ in individual cells injected with increasing doses of Rad RNA. N=1 experiment; statistics: paired t-test. **E**, Fold change increase in I_Ba_ caused by Iso as a function of Rad:β_2b_ RNA ratio. N=1 experiment. Statistics: One Way ANOVA.

**Figure 6.**
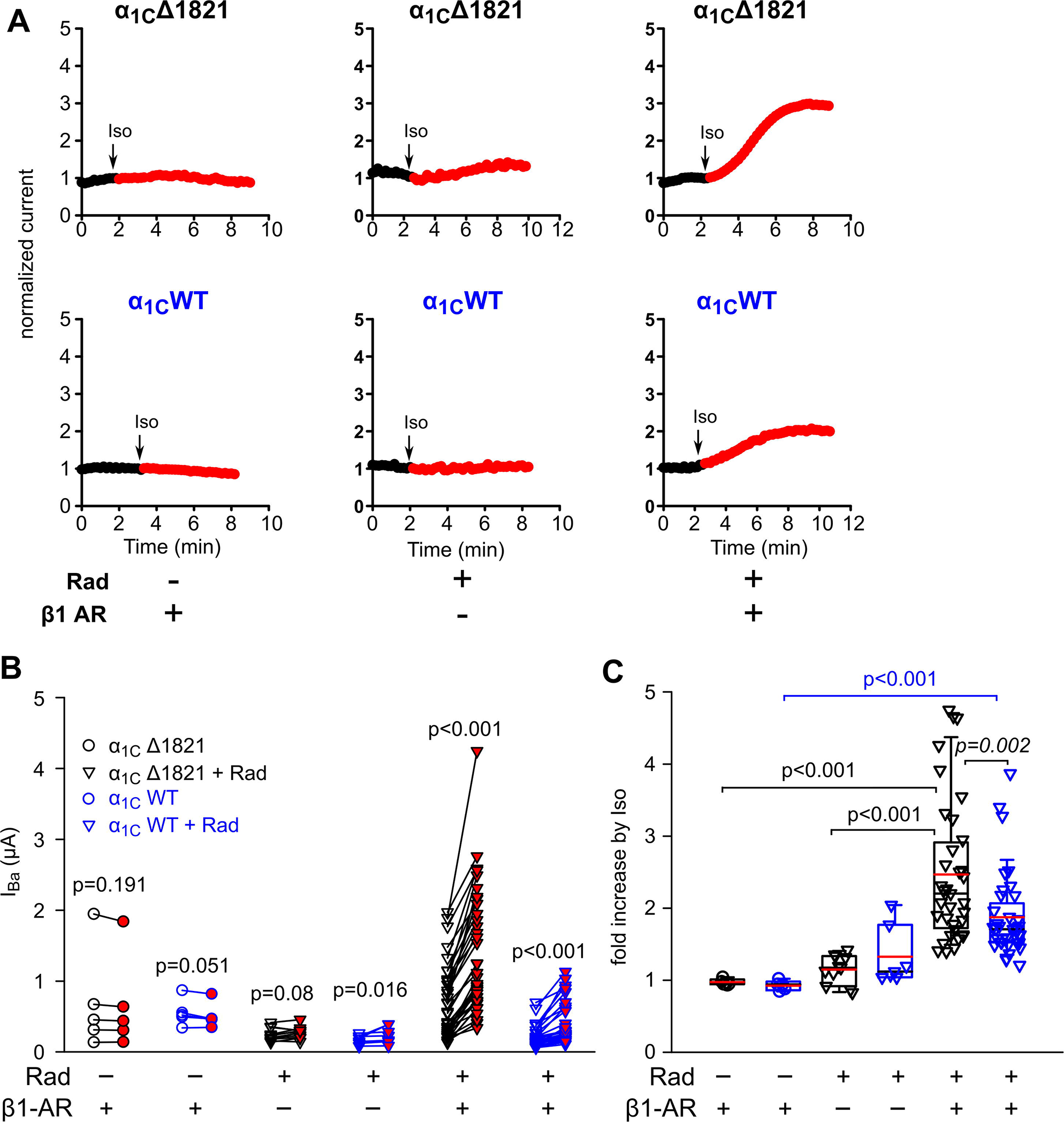
β1-AR regulation of full-length and truncated α_1C_. **A**, Representative diary plots of I_Ba_ in oocytes expressing wt (full-length) α_1C_ (Ca_V_1.2-α_1C_) or Ca_V_1.2-α_1C_Δ1821 channels (lower and upper panels, respectively) with or without Rad and β1-AR, and the response to 50 μM Iso. β_2b_ and α2δ were coexpressed in all cases. **B**, “Before-after” plots of Iso-induced changes in I_Ba_ in individual cells. The Rad:β2b ratio in oocytes expressing wt Ca_V_1.2 (blue symbols) or Ca_V_1.2Δ1821 (black symbols) was 1:3 and 1:2, respectively. Lower ratio was used in cells expressing wt-α_1C_ to allow higher basal currents. N=3; statistics: paired t-test. **C**, Rad is essential for the Iso-induced increase in I_Ba_. N=3 experiments. Statistical analysis was performed separately for wt Ca_V_1.2-α_1C_ and Ca_V_1.2-α_1C_Δ1821 with ANOVA on ranks (p<.001). In addition, Mann-Whitney Rank Sum test was used to compare the last two groups, wt Ca_V_1.2 vs. Ca_V_1.2Δ1821 with Rad and β1-AR (p=0.002). For wt Ca_V_1.2 with Rad and β1-AR, there was no significant difference from the group without β1-AR but with Rad (p=0.076, One-Way ANOVA on ranks, Dunnett’s test).

In cardiomyocytes, both full-length (wt) and truncated α_1C_ are present, but it is not known if both isoforms are equally regulated by β1-AR. To study β1-AR regulation of Ca_V_1.2 containing wt-α_1C_, we first titrated Rad:β_2b_, in oocytes that coexpressed β1-AR. As for α_1C_Δ1821 (see Fig. 1B), we found an inverse correlation between Rad concentration and I_Ba_ (r=-0.85, p=0.002; Fig. S3). In addition, compared with α_1C_Δ1821, lower Rad:β_2b_ RNA ratios were sufficient to yield a significant increase in currents upon perfusion of Iso (Fig. 5D-E, compare with Fig. 1E). In contrast, in the absence of Rad, there was no increase in currents over time during Iso perfusion (Fig. 5D-E; Fig. 6). Based on the results of Rad titration of Fig. 5, we used Rad:β_2b_ RNA ratio of 1:3 to 1:2 thereafter for channels containing wt-α_1C_.

We next systematically compared the effect of Iso on channels containing either wt-α_1C_ or α_1C_Δ1821, with or without coexpressed β1-AR and Rad (Fig. 6). Without the coexpression of Rad, activation of β1-AR did not produce any increase in I_Ba_ in either full-length or truncated channel. This result indicates that the Rad-independent pathway is not activated by β1-AR under the conditions used. Moreover, it appears that oocytes do not contain endogenous Rad or similar RGK proteins that are available for the β1-AR – Ca_V_1.2 cascade. Interestingly, when oocytes expressed Rad without the receptor, Iso caused a small increase in I_Ba_: ~15% in α_1C_Δ1821 (which did not reach statistical significance, p=0.08 by paired t-test), and ~33% in wt-α (p=0.016) (Fig. 6B, C). These results corroborate a previous report^40^ suggesting that endogenous β-AR is present in some oocyte batches.

In oocytes that expressed β1-AR, Rad and Ca_V_1.2, Iso induced a robust increase in I_Ba_ (Fig. 6A, B; p<.001 for both α_1C_ forms). However, comparison of Iso-induced increase in the two α_1C_ forms revealed a significantly greater effect on the truncated channel than on the full-length α_1C_ (147% vs. 87% increase in mean I_Ba_, respectively; median fold increase 2.2 [IQR 1.72-2.91] versus 1.71 [IQR 1.53-2.07], p=0.002; Fig. 6C). These observations are in agreement with the greater effect of cAMP on the truncated channel (Fig. 4) and imply that the dCT does play a role in attenuating the PKA-induced augmentation of Ca_V_1.2 currents, at least when it is not cleaved from α_1C_.

### *Reconstitution of the* β*2 adrenergic receptor regulation of Ca*_*V*_*1.2*

In healthy heart, the distribution of β2-AR is limited to specific parts of the heart (atrium, apex) and mainly to T-tubules within cardiomyocytes, but becomes more widely distributed over the cardiomyocyte surface in failing heart^11, 41, 42^. β2-AR is also the major β-AR form in the nervous system^9^. We began with titration of β2-AR with Ca_V_1.2 containing α_1C_Δ1821. We expressed the receptors by injecting their RNAs at 50 and 200 pg RNA/oocyte. We also injected higher RNA doses, but above 1 ng/oocyte, cells showed low rate of survival during incubation, and surviving oocytes had high leak currents.

As shown before with cAMP injection (Fig. 1), in oocytes expressing Ca_V_1.2-α_1C_Δ1821, Rad and β1-AR, a robust regulation of I_Ba_ by Iso was observed (Fig. 7A upper trace, Fig. 7 B, C). Surprisingly, in oocytes of the same batch expressing Ca_V_1.2-α_1C_Δ1821, β2-AR and Rad, I_Ba_ did not respond to Iso (Fig. 7A lower panel, Fig. 7B-C). In some cases, we did observe an increase in I_Ba_ (e.g. Fig. 8G) but it was very small compared to β1-induced Ca_V_1.2 stimulation. To test if the expressed β2-AR was functioning well, we used cystic fibrosis transmembrane conductance regulator (CFTR) channel as a control. PKA phosphorylation of CFTR, a chloride channel, activates the channel, leading to an increased outward chloride current^43^. Moreover, CFTR is robustly activated by cAMP and PKA-CS injection in *Xenopus* oocytes^44^.

**Figure 7.**
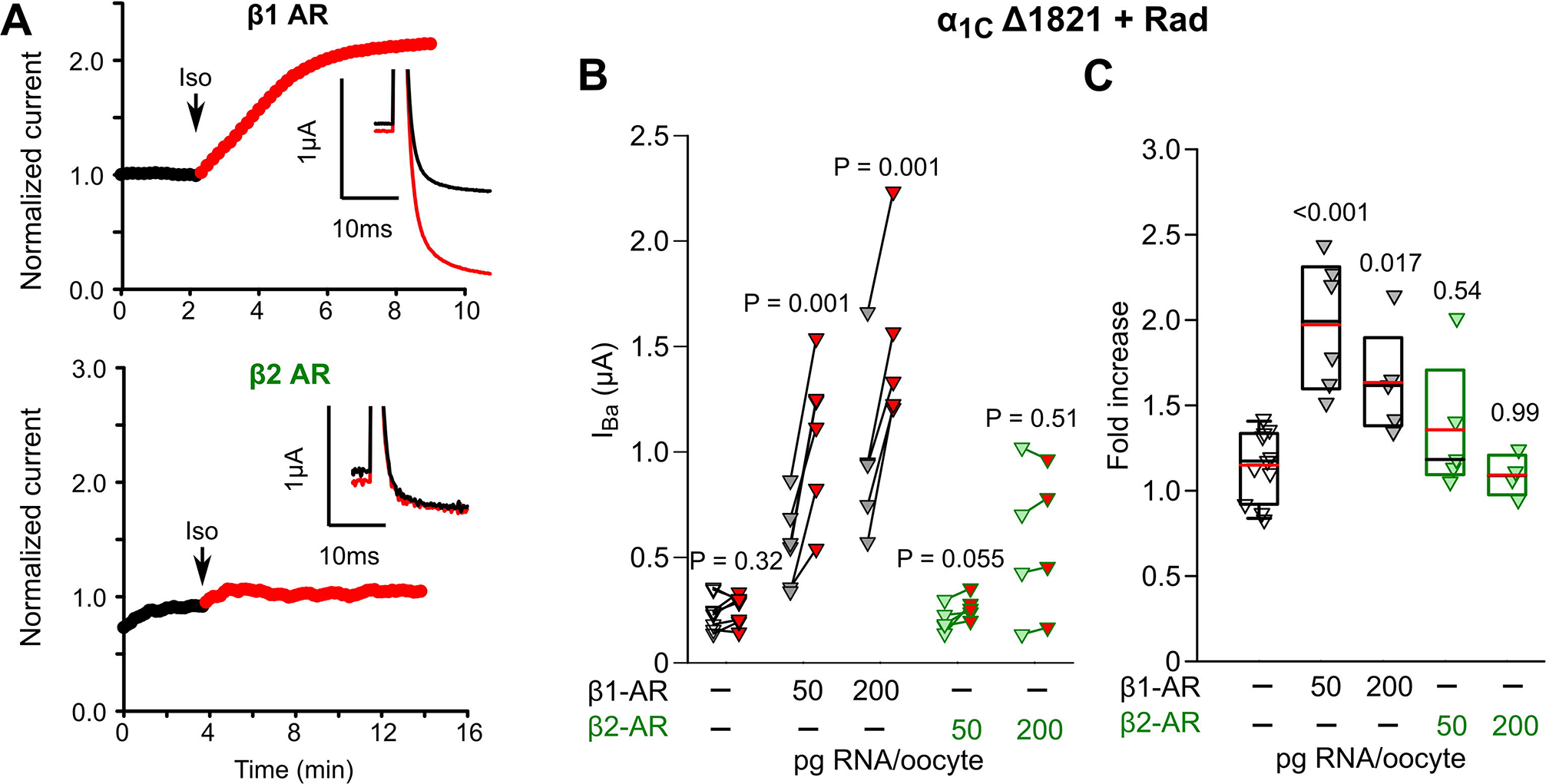
Comparison of β1 and β2 adrenergic regulation of Ca_V_1.2. **A**, Representative diary plots of I_Ba_ showing the effect of Iso on Ca_V_1.2-α_1C_Δ1821 in oocytes co-expressing β1-AR (upper panel) or β2-AR (lower panel). Insets show I_Ba_ at +20 mV before and 9-10 min after the addition of Iso. RNAs of both receptors were made on the template of cDNAs inserted into the pGEM-HJ vector. **B**, “Before-after” plots of Iso-induced changes in I_Ba_ in individual cells co-expressing Ca_V_1.2-α_1C_Δ1821 and Rad, without any receptor or with either β1-AR or β2-AR. Data are shown for receptors RNAs concentrations of 0, 50 and 200 pg RNA/oocyte. Rad:β_2b_ RNA ratio was 1:2. N=1 experiment; statistics: paired t-test. **C**, Fold change in I_Ba_ induced by Iso (summary of data shown in A and B). Statistics: One Way ANOVA followed by Dunnett’s test.

**Figure 8.**
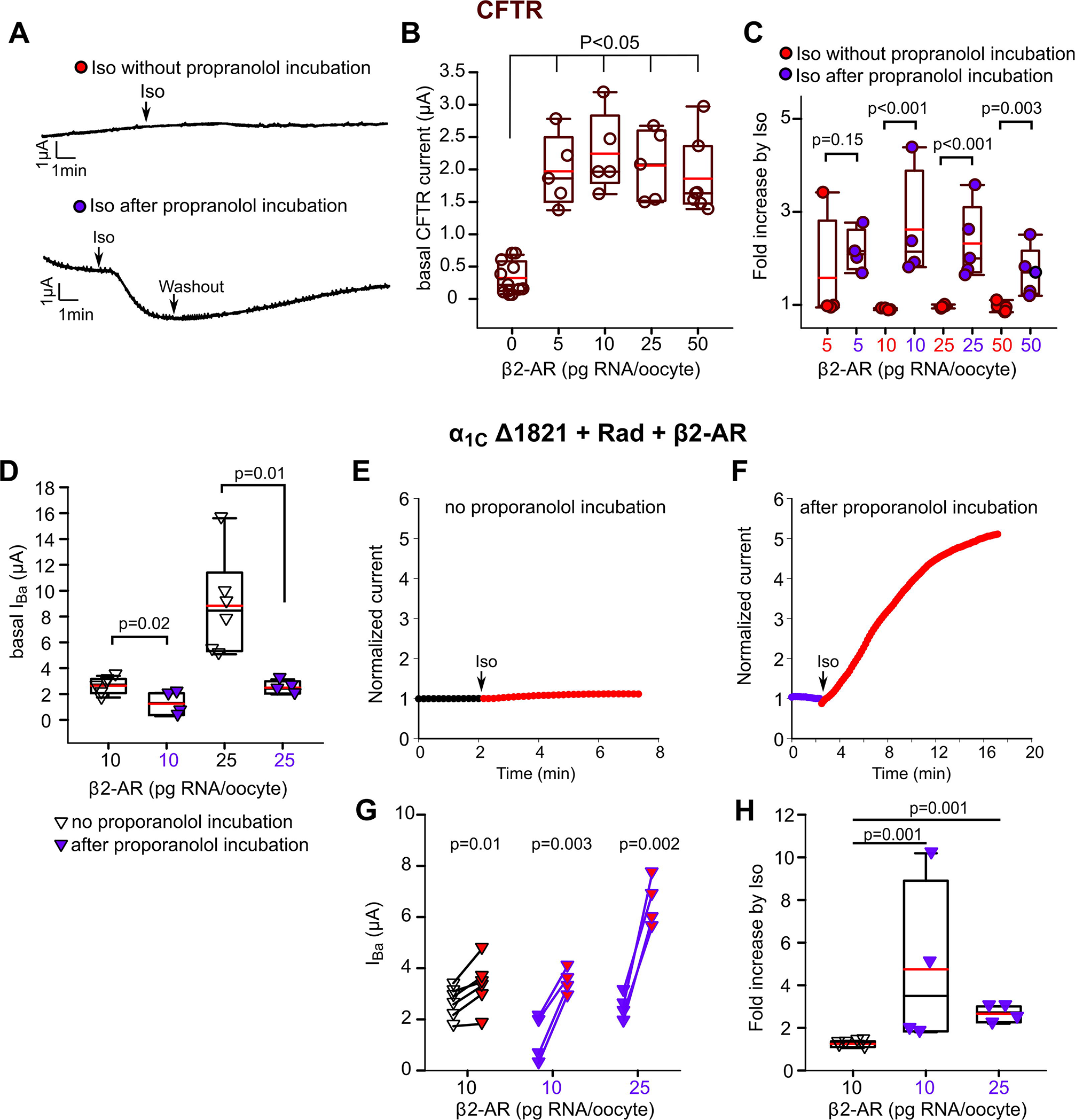
Reconstitution of the β2 adrenergic receptor regulation of CFTR and α_1C_. **A**, Preincubation with propranolol is needed to reinstate the adrenergic regulation of CFTR by β2-AR. Oocytes expressed CFTR and β2-AR (1 and 0.005-0.05 ng/oocyte, respectively). CFTR chloride currents were recorded in gap free mode at −80 mV. 50 μm Iso was applied after stabilization of the current (about 2 min after start of record). Upper panel shows a representative record without propranolol incubation. The lower panel shows a representative oocyte preincubated in 10 μM propranolol for 60-120 min, then transferred to the experimental chamber, voltage clamped (start of record) and perfused with propranolol-free solution for about 2 min prior to the addition of Iso. **B**, Basal chloride currents in oocytes co-expressing CFTR channel (1 ng RNA/oocyte) and β2-AR (increasing doses of RNA). Oocytes expressing CFTR alone had statistically significant lower basal currents than oocytes co-expressing β2-AR (N=1 experiment; statistics: p<.001, Kruskal-Wallis One Way ANOVA on Ranks) **C**, Fold change in chloride current by Iso. The plot compares the effects of 50 μM Iso in oocytes injected with the indicated doses of β2-AR RNA, without (red symbols) or with (purple symbols) propranolol preincubation. N=1 experiment. Statistics: Mann-Whitney Rank Sum Test. **D**, Comparison of basal I_Ba_ in oocytes co-expressing Ca_V_1.2-α_1C_Δ1821, Rad and β2-AR, which were incubated without propranolol (black) or with 10 μM propranolol (blue). I_Ba_ was recorded at peak current of +20 mV. Oocytes were preincubated with 10 μM propranolol for 60-120 min, then transferred to the experimental chamber, voltage clamped to +20 mV and perfused with propranolol-free 40 mM Ba^2+^ solution until stabilization of the current for 2 min. Propranolol washout lasted approximately 1-2 minutes. Oocytes co-expressed α_1C_Δ1821, β_2b_, α2δ, Rad (Rad:β2_b_ RNA ratio was 0.5). N=1; statistics: t-test. **E-F**, Representative diary plots of oocytes co-expressing α_1C_Δ1821, Rad and β2-AR without (left) and with (right) preincubation with propranolol. **G**, “Before-after” plots of Iso-induced changes in I_Ba_ in individual cells. Blue symbols represent data form oocytes preincubated with propranolol, black – without preincubation. N=1; statistics: paired t-test. **H**, Fold increase in I_Ba_ induced by Iso (summary of data shown in G). Oocytes preincubated with propranolol (blue) had statistically significant higher fold increase. N=1; statistics: Kruskal-Wallis One Way ANOVA on Ranks followed by Dunnett’s test.

Co-expressing CFTR with a range of β2-AR doses (RNA range, 5-50 pg/oocyte) was associated with significantly higher basal chloride currents (I_CFTR_) compared with cells expressing CFTR alone (p<.001) (Fig. 8B). Iso perfusion did not cause an increase in I_CFTR_ at any of the β2-AR doses (Fig. 8A upper panel, Fig. 8C, red circles). This observation implied that CFTR channels may have already been opened by agonist-independent constitutively active β2-ARs, thus blunting any response to Iso. Reducing the basal activity of the receptor would result in lower chloride basal currents and may reinstate the activation of β2-AR upon Iso perfusion. Propranolol is a non-selective β blocker and an inverse agonist, which is known to reduce the constitutive activity of β2-AR^45^. Therefore, we incubated the oocytes for 1-2 hours with 10 μM propranolol before starting the recording. Just before the recording, the oocyte was placed in the experimental chamber, voltage clamp was established, and the cell was washed with propranolol-free solution for 2-4 minutes before application of Iso. With propranolol preincubation, we found that Iso induced a robust 1.5-3 fold increase in chloride currents in oocytes expressing CFTR and β2-AR (Fig. 8A lower panel and Fig. 8C, purple circles).

The results of the CFTR experiment strongly supported the possibility that high constitutive activity of β2-AR precluded further effect of Iso, also in the case of Ca_V_1.2. Therefore, we employed the propranolol preincubation protocol for Ca_V_1.2-α_1C_Δ1821, coexpressed with β2-AR and Rad. We measured I_Ba_ in oocytes not preincubated (black) or pre-incubated (purple) with propranolol (Fig. 8D). As predicted, preincubation with propranolol resulted in a statistically significant reduction of basal I_Ba_ (Fig. 8D). Application of Iso produced only a mild but significant increase in I_Ba_ without propranolol incubation, by 24±5%, in this experiment (Fig. 8E, G, H) compared to a significantly greater increase, 3-5 fold, with propranolol pre-incubation (Fig. 8F-H). We conclude that we successfully reconstituted adrenergic regulation of the Ca_V_1.2 channel current as mediated by the two canonical and physiologically-relevant adrenergic receptors.

## Discussion

Numerous biological regulations of ion channels that have been fully reconstituted in heterologous systems accelerated the understanding of their mechanisms and structure-function relationships. However, the classical adrenergic regulation of cardiac L-type Ca^2+^ channel remained an unmet challenge for several decades. The recent discovery of the crucial role of Rad^15^ was a turning point. Here we report, for the first time, the heterologous reconstitution of the full cascade of β-AR regulation of the cardiac L-type Ca^2+^ channel, Ca_V_1.2, in *Xenopus* oocytes, starting with the receptor. Two major advantages of the *Xenopus* oocyte model are accurate titration of protein expression (by titrated RNA injection), and the ability to co-express a large number of proteins. We utilized the simplicity and robustness of the oocyte expression system to reconstitute the full β-AR cascade, to address the role of Rad, the relation between Rad-dependent and the previously reported Rad-independent PKA regulation^24^, and to elaborate the role of Ca_V_β and the distal parts of N- and C-termini of α_1C_.

First, we confirmed the cAMP-induced upregulation of I_Ba_ in oocytes expressing Ca_V_1.2-αΔ1821 without Rad^24^ which, in this series of experiments, was ~20%. This is less than the previously reported 30-40%^24^, probably because here we injected about half the amount of cAMP. Notably, a greater Rad-independent I_Ba_ potentiation was attained by injecting purified PKA-CS (Fig. S1).

Next, we scrutinized the role of Rad, initially using cAMP injection to activate the oocyte’s endogenous PKA. Liu et al. reported a ~1.5-2-fold increase in maximal conductance (G_max_) of Ca_V_1.2 containing the full-length α_1C_ in HEK cells^15^. Here, with α_1C_Δ1821, titrated expression of Rad showed the expected^26, 27^ Rad-dose-dependent decrease in I_Ba_, paralleled by a robust enhancement in cAMP-induced increase in I_Ba_, up to ~2.2-fold (Fig. 1). These results confirm the importance of Rad in PKA regulation and, moreover, suggest that Rad-dependent regulation applies to both full-length and C-terminally-truncated α_1C_. Importantly, the Rad-induced enhancement by cAMP leveled off and occasionally seemed to diminish at higher Rad:Ca_V_1.2 RNA ratios (we expressed all channel subunits in equal weights doses), suggesting that molar excess of Rad may override the PKA-induced abolition of tonic Rad inhibition. Thus, we used the optimal Rad:β_2b_ RNA ratio, 1:3 – 1:1, in all experiments.

We next inquired whether Rad-dependent and Rad-independent regulatory mechanisms are distinct, considering the possibility that the latter was actually the same as Rad-dependent regulation, possibly mediated by an endogenous RGK protein in the oocyte. Our results unequivocally demonstrate that the two modalities are carried out by distinct molecular mechanisms. First, Rad-dependent regulation was fully Ca_V_β-dependent, as demonstrated^15^. Both Rad inhibition of the basal I_Ba_, and the Rad-dependent enhancement of cAMP effect critically depended on the presence of coexpressed Ca_V_β subunit (full-length or core), and both Rad actions were suppressed by a triple mutation that abolishes Rad-Ca_V_β interaction (Fig. 2). In contrast, the Rad-independent cAMP effect was the same with all β_2b_ constructs used, or without coexpressed Ca_V_β. Second, as shown before^24^, the Rad-independent regulation crucially depended on two cytosolic elements of α_1C_: the inhibitory module (initial segment) of the NT (Fig. 3, Fig. S1), and truncation of dCT of α_1C_ (Fig. 4). In contrast, in the presence of Rad, robust regulation of I_Ba_ by cAMP was consistently observed, both after the deletion of the initial NT segment, and in channels containing either full-length or dCT-truncated α_1C_.

We then reconstituted the full β-AR cascade with coexpressed β1-AR. Activation of β1-AR by Iso caused a >2-fold increase in Ca_V_1.2-α_1C_Δ1821 current and the typical hyperpolarizing shift in voltage dependence of activation (Fig. 5, 6), resembling cardiomyocytes^16^. Coexpression of G_s_ and adenylyl cyclase was not necessary, suggesting sufficient levels of endogenous proteins. However, expression of Rad was essential; in its absence, no increase in I_Ba_ was observed. It is unclear why the Rad-independent regulation of Ca_V_1.2-α_1C_Δ1821 could not be produced by activation of β1-AR. We consider several possibilities, among them another missing factor needed for this regulation, or a stoichiometry predicament with either the receptor or a downstream protein of the cascade. Unfortunately, expressing high doses or β1-AR, Gα_s_ or adenylyl cyclase consistently resulted in high oocyte mortality. Whatever the reason, it appeared that, under the conditions of our experiments, β1-AR regulated Ca_V_1.2 only via the Rad-dependent mechanism.

Of particular importance is the observation that, once Rad was present, both dCT-truncated and full length channels were upregulated both by cAMP and by β1-AR (Figs. 5, 6). This was never clear, and previously controversial^18^. The full-length α is present in the heart and seems even more abundant in neurons^9^. Yet, the mechanism of regulation of neuronal Ca_V_1.2 appears different from that of cardiac; the direct PKA phosphorylation of serine 1928 (located in the dCT) is highly important in neurons^46^, but not in the heart^13, 47^. However, in our system the β1-AR activation of full-length α_1C_-based channels (containing S1928) still required Rad. This indicates that phosphorylation of S1928 alone is not enough. Interestingly, a more detailed examination of the results reveals a potentially important difference: with cAMP, and even more so with β1-AR, the overall regulation of the dCT-truncated channel is significantly stronger than of the full-length channel. A more straightforward interpretation would be the contribution of Rad-independent regulation, which is present only in α_1C_Δ1821. However, this is valid only for cAMP-induced regulation; as discussed, β1-AR does not appear to initiate the Rad-independent pathway. We propose that the dCT exerts a regulatory control over the Rad-dependent β-AR regulation of the channel, via mechanisms that need to be explored.

Finally, we were also able to reconstitute the Ca_V_1.2 enhancement by β2-AR which, like β1-AR, acted via the Rad-dependent pathway (Figs. 7, 8). However, there was a significant difference. As demonstrated with both CFTR and Ca_V_1.2, β2-AR constitutively activated the G_s_-PKA pathway, rendering very high basal CFTR and Ca_V_1.2 currents, as revealed by the inverse agonist, propranolol (Fig. 8). The basal Ca_V_1.2 activation was so strong even with very low doses of β2-AR RNA, that pretreatment with propranolol was required to observe upregulation by Iso. Whereas high basal constitutive activity of β2-AR is well established^45, 48, 49^, in the heart, agonist stimulation of β2-AR normally increases Ca_V_1.2 currents^39, 42^. We assume that, in cardiomyocytes, specific mechanisms such as restricted localization^9, 41, 42^ or additional auxiliary proteins may regulate the basal activity of β2-AR.

In summary, we reconstituted the β1-AR and β2-AR regulation of Ca_V_1.2 in the *Xenopus* oocyte model system. We demonstrate a robust (~2-fold) enhancement of Ca^2+^ channel currents, by cAMP and Iso, in a Rad-dependent manner, and confirm the existence of a less prominent, separate, Rad-independent PKA regulation of Ca_V_1.2 (~30% increase in Ca_V_1.2 current). Our reconstituted system replicates the known basic features of β-AR regulation of cardiac Ca_V_1.2 and reveals novel molecular details, including the consequences of proteolytic processing of α_1C_ with respect to β-AR regulation, the differential roles of Ca_V_β subunit and of the N- and C-termini of α_1C_ in the Rad-dependent and Rad-independent regulation, and the differences in channel regulation by β1-AR and β2-AR. The reconstitution of the basic cascade will enable further investigation of the mechanisms of Rad action and additional auxiliary proteins implicated in macromolecular complexes involved in β-AR regulation of Ca_V_1.2, such as A-kinase anchoring proteins, phosphatases and phosphodiesterases, and others^50^. It may also be instrumental in identifying potential targets for therapeutic modulation of β-adrenergic regulation in heart and other tissues.

