## Supplemental Materials for "Reconstitution of β-adrenergic regulation of Ca_V_1.2: Rad-dependent and Rad-independent protein kinase A mechanisms"

### Supplemental Material

#### Expanded Methods

##### *Experimental animals and ethical approval*

Experiments were approved by Tel Aviv University Institutional Animal Care and Use Committee (permits # 01-16-104 and 01-20-083). Adult female *Xenopus laevis* frogs were purchased from Xenopus-1 (Dexter, MI, USA). The frogs were housed in Tel Aviv University Medical School Animal Facility and handled essentially as described[24]. Female frogs were maintained at 20±2°C on 10 h light/14 h dark cycle. Frogs were anaesthetized in 0.2% tricaine methanesulfonate (MS-222), and portions of ovary were removed through an incision on the abdomen. The incision was sutured, and the animal was held in a separate tank until it had fully recovered from the anesthesia, and afterwards was returned to a separate tank for post-operational animals. The animals did not show any signs of post-operational distress and were allowed to recover for at least 3 months until the next surgery. Following the final collection of oocytes, anaesthetized frogs were sacrificed by decapitation and double pithing.

##### *DNA constructs and RNA*

The cDNA constructs used are: Human Rad (NP_001122322), cardiac long-N terminus (LNT) isoform of rabbit $\alpha_{1C}$ (GenBank: X15539) and the corresponding mouse $\alpha_{1C}$ isoform (NM_001255999.2), rabbit $Ca_V\beta_{2b}$ (originally termed $\beta_{2a}$[29]; GenBank: X64297.1; $Ca_V\beta2N4$ according to the comprehensive nomenclature[4]), $\alpha_2\delta_1$ (GenBank: M21948). All DNA mutations have been introduced using standard PCR-based procedures and verified by sequencing of the coding region of the cDNA. cDNAs of rabbit and mouse $\alpha_{1C}\Delta821$, in which distal C terminus is truncated, were prepared by the removal of the cDNA segment encoding the dCT from the plasmid and introducing a stop codon after valine 1821. Rabbit $\alpha_{1C}\Delta20\Delta1821$ was prepared by deletion of the first 20 amino acids of the LNT of $\alpha_{1C}\Delta1821$. Two NT mutants of mouse $\alpha_{1C}$ were constructed: $\alpha_{1C}$NT-4A$\Delta$1821, with alanine substitution of the first 4 amino acids; and $\alpha_{1C}$NT-TYP$\Delta$1821, with alanine substitution of amino acids $T_{10}$, $Y_{13}$ and $P_{15}$. All mouse $\alpha_{1C}$ constructs were inserted into pMXT vector, and contained the double mutation T1066Y/Q1070M which renders the channel dihydropyridine-insensitive, but does not affect regulation by PKA[14]. The $\beta_{2b}$-core construct was prepared by deletion of the first 25 amino acids in the N-terminus of $Ca_V\beta_{2b}$, deletion of amino acids 423-606 in the C-terminus and removing the linker amino acids 138-202 by PCR[30]. $\beta_{2b}$-core$_{3Da}$ was prepared similarly, but the first 25 amino acids were not deleted and 3 point mutations (D244A/D320A/D322A ) were introduced by PCR using standard site-directed mutagenesis (residues numbering according to Opatowsky et al.[30]). Human CFTR channel (NM_00492) was in pSP64 vector; we note that this cDNA had an extra serine in position 264, which did not affect the channel's function. Human β2 adrenergic receptor (GenBank: AAA88015.1) was subcloned into pGEM-HJ vector. Mouse β1 adrenergic receptor (NP_031445.2) was in pcDNA3.1 vector. For comparison with β2 adrenergic receptor, it was subcloned into pGEM-HJ vector. All other cDNA constructs used for RNA synthesis were inserted into pGEM-GSB or pGEM-HJ vectors, all of which contain 5' and 3' UTR from *Xenopus* β-globin.

The RNAs were prepared using a standard procedure[24]. $Ca_V1.2$ was expressed by injecting the RNAs of all subunits ($\alpha_{1C}$, $\beta_{2b}$, $\alpha_2\delta1$) in equal amounts, unless indicated otherwise. The amount of injected RNA, per oocyte, for full-length $\alpha_{1C}$ was 1.8-5 ng. For $\alpha_{1C}\Delta1821$

constructs, the injected RNA amounts were: 1-1.5 ng for $\alpha_{1C}\Delta1821$, $\alpha_{1C}\Delta20\Delta1821$ 0.6 ng, $\alpha_{1C}$NT-4AΔ1821 and $\alpha_{1C}$NT-TYPΔ1821 3 ng. When $Ca_V\beta$ was not expressed, we injected 5 ng RNA of $\alpha_{1C}\Delta1821$ and $\alpha_2\delta1$. Rad RNA was injected in Rad:$\beta_{2b}$ ratio of 1:3-1:1. RNA of β1-AR was injected at 5 ng/oocyte if produced on the template of pcDNA vector, or 0.05-0.2 ng when the DNA was cloned into pGEM-HJ. β2-AR was in pGEM-HJ and 0.05-0.2 ng/oocytes were injected. For titration purpose we injected even higher doses (0.5-5 ng), however those doses decreased the viability of the oocytes and led to fast deterioration in leak current after insertion of the recording electrodes.

##### *Purification of recombinant PKA-CS*

We used His-tagged catalytic-subunit of PKA (His-PKA-CS, GenBank: NM 008854.5) for protein purification from *E. coli*. His-PKA-CS protein was expressed in the *E.coli* strain BL21. Cells were grown in YT medium containing 100 μg/ml ampicillin and 36 μg/ml chloramphenicol at 37$^{o}$ C to an optical density at 600 nm of 0.5-0.8, induced with 1 mM IPTG and grown for an additional 12h at 16$^{o}$ C, collected by centrifugation, and stored frozen. Cells from 1500 ml of culture were resuspended in 40-ml lysis buffer (50 mM sodium phosphate, 100 mM NaCl, 20 mM Tris-HCl, 5 mM β mercaptoethanol (β-ME), pH 8.0) and protease-inhibitor cocktail tablet (Roche), 2 mM phenylmethylsulfonyl fluoride (PMSF) and lysed in a microfluidizer. Thereafter, cells were centrifuged at 17,000 rpm at 4$^{o}$C for 30 minutes. The lysate was applied to the Ni-NTA agarose column (2 ml/min). Samples were taken from lysate and flow-through. Then washing was carried out in wash buffer: 50 mM $KH_2PO_4$, 20 mM Tris–HCl (pH 8), 100 mM NaCl, 10 mM imidazole, 5 mM β-ME. Then elution step was carried out with 40 ml of elution buffer: 250 mM imidazole, 50 mM $KH_2PO_4$, 20 mM Tris–HCl (pH 8), 100 mM NaCl+ 5 mM β-ME. The eluate was concentrated with Centricon 10 kDa to 5 ml and added to 80 ml gel-filtration buffer (20 mM Mops (pH 8), 100 mM KCl, 5 mM β-ME) and purified on a Superdex G-75 column, subdivided into 4-5 µl aliquots (5µg/µl) and stored at -80°C.

##### *Electrophysiology*

Oocytes were defolliculated by collagenase, injected with RNA and incubated for 3 days before recording at 20–22°C in NDE solution (in mM: 96 NaCl, 2 KCl, 1 $MgCl_2$, 1 $CaCl_2$, 5 Hepes, 2.5 pyruvic acid, and 0.1 gentamycin sulfate)[24]. Ion channel currents in oocytes were measured using the two-electrode voltage clamp technique with a GeneClamp 500 amplifier (Molecular Devices, Sunnyvale, CA, USA). CFTR currents were measured at −80 mV in ND96 solution (in mM: 96 NaCl, 2 KCl,1 $MgCl_2$, 1 $CaCl_2$, 5 Hepes, pH 7.6). Whole-cell $Ba^{2+}$ current ($I_{Ba}$) was elicited by 20 ms depolarizing pulses from a resting potential of −80 mV to 20 mV, with a 10 s interval between sweeps, in 40 mM $Ba^{2+}$ solution (in mM: 40 $Ba(OH)_2$,50 NaOH, 2 KOH, and 5 Hepes, titrated to pH 7.5 with methanesulfonic acid). In one experiment, i.e., **Fig. 3**, for $\alpha_{1C}$Δ20Δ1821, $I_{Ba}$ amplitudes in 40 mM $Ba^{2+}$ solution exceeded 6 µA. In this case, to avoid artifacts resulting from oocyte series resistance, recordings were made in 2 mM $Ba^{2+}$ solution (in mM: 2 $Ba(OH)_2$, 96 NaOH, 2 KOH, and 5 Hepes, titrated to pH 7.5 with methanesulfonic acid). These measurements were used to assess cAMP-induced changes in $I_{Ba}$ amplitude only and have not been included in amplitude summaries and activation curve fits.

In the current–voltage (I–V) protocols, currents were elicited by 20 ms pulses from the holding potential −80 mV to voltages from −70 to +80 mV with 10 mV intervals and 10 s between sweeps. Currents measured in the presence of 200 µM $Cd^{2+}$ were subtracted from total $I_{Ba}$ (**Fig. 5A-C***)* to yield the net $I_{Ba}$. I-V curves were analyzed as described[32]. In each cell, I-V curves in the range -70 to +40 mV were fitted to the Boltzmann equation in the form:

$I = G_{max}(V_m - V_{rev})/(1 + exp(-(V_m - V_{1/2})/K_a))$,

where $G_{max}$ is the maximal $Ba^{2+}$ conductance, $V_m$ is the membrane voltage, $V_{rev}$ is the reversal potential of the current, $K_a$ is the slope factor and $V_{1/2}$ is half-maximum activation voltage. The parameters obtained for $G_{max}$ and $V_{rev}$ were then used to calculate fractional conductance data points at each $V_m$ using the equation:

$G/G_{max} = I/(G_{max}(V_m - V_{rev}))$.

Conductance-voltage (G–V) curves through the data points were plotted with the values of $V_{1/2}$ and $K_a$ obtained from the fit of the I-V curves, using the following form of the Boltzmann equation:

$G/G_{max} = 1/(1+exp(-V_m-V_{1/2})/K_a))$.

##### *Injection of cAMP and PKA-CS and Isoproterenol perfusion*

cAMP (Sigma, A6885) was diluted in $H_2O$, stored in small aliquots as 40 mM stock solution at -20$^o$C, and thawed only once. For injection, cAMP was diluted to 20 mM. Purified His-PKA-CS was kept in aliquots of 5 µg/µl and stored in small aliquots at -80$^O$C, thawed at the beginning of the experiment and during the experiment kept on ice. The micropipette tip was trimmed so that application of pressure during 1-2 s extruded approximately 5 nl (0.5% of oocyte volume). Injection during recording was done with sharp capillary glass micropipettes filled with cAMP or PKA-CS. The injection needle was inserted after the whole-cell voltage clamp has been established. Compounds were injected into oocytes with pressure only after observing that currents have been stable for at least 2 minutes. The concentration of injected cAMP in the oocytes was ~100 µM, assuming oocyte volume of 1 µl. The final amount of PKA-CS injected was ~25 ng/oocyte. Injection artifacts visible as a sharp shift in current, accompanied by an increase in leak current, were usually minor (see **Figs. 1C, 2B**). Records with injection artifact that exceeded 10% of $I_{Ba}$ amplitude were discarded.

Isoprenaline hydrochloride (isoproterenol, Sigma-Aldrich, I5627) was diluted in $H_2O$ and kept in 100 mM stock solution aliquots. Perfusion of isoproterenol 50 µM began after documentation of a stable peak current for 2 minutes. Propranolol (Sigma-Aldrich, P0844) was in dissolved DMSO at 50 mM. For the propranolol pretreatment procedure, oocytes were incubated for 1-2 hours in 10 µM propranolol in NDE solution.

***Statistical analysis***

In all experiments, the fold change in current caused by an externally applied or injected substance was calculated by dividing the current at the end of the recording by the current measured before substance application in the same cell. The values before and after treatment with cAMP/PKA-CS/Isoproterenol were compared using paired t-test for normally distributed variables, otherwise a Wilcoxon test was performed. Two-sample comparisons of different treatment groups were done by independent sample t-test, or by Mann-Whitney test for data that did not pass normality test. Comparison of multiple test groups was done with One Way ANOVA if the data was normally distributed or Kruskal-Wallis ANOVA on ranks when the data did not distribute normally. A Bonferroni post hoc test was performed for normally distributed data and Dunnett's post hoc test otherwise.  Statistical analysis was performed with SigmaPlot 13 (Systat Software Inc., San Jose, CA, USA). Data sets that did not pass the Shapiro-Wilk normality test were reported as median and interquartile range (IQR) [Q1-Q3]. Normally distributed continuous variables were reported as mean±standard error of mean.

#### Supplemental Figures.

***Fig. S1.***

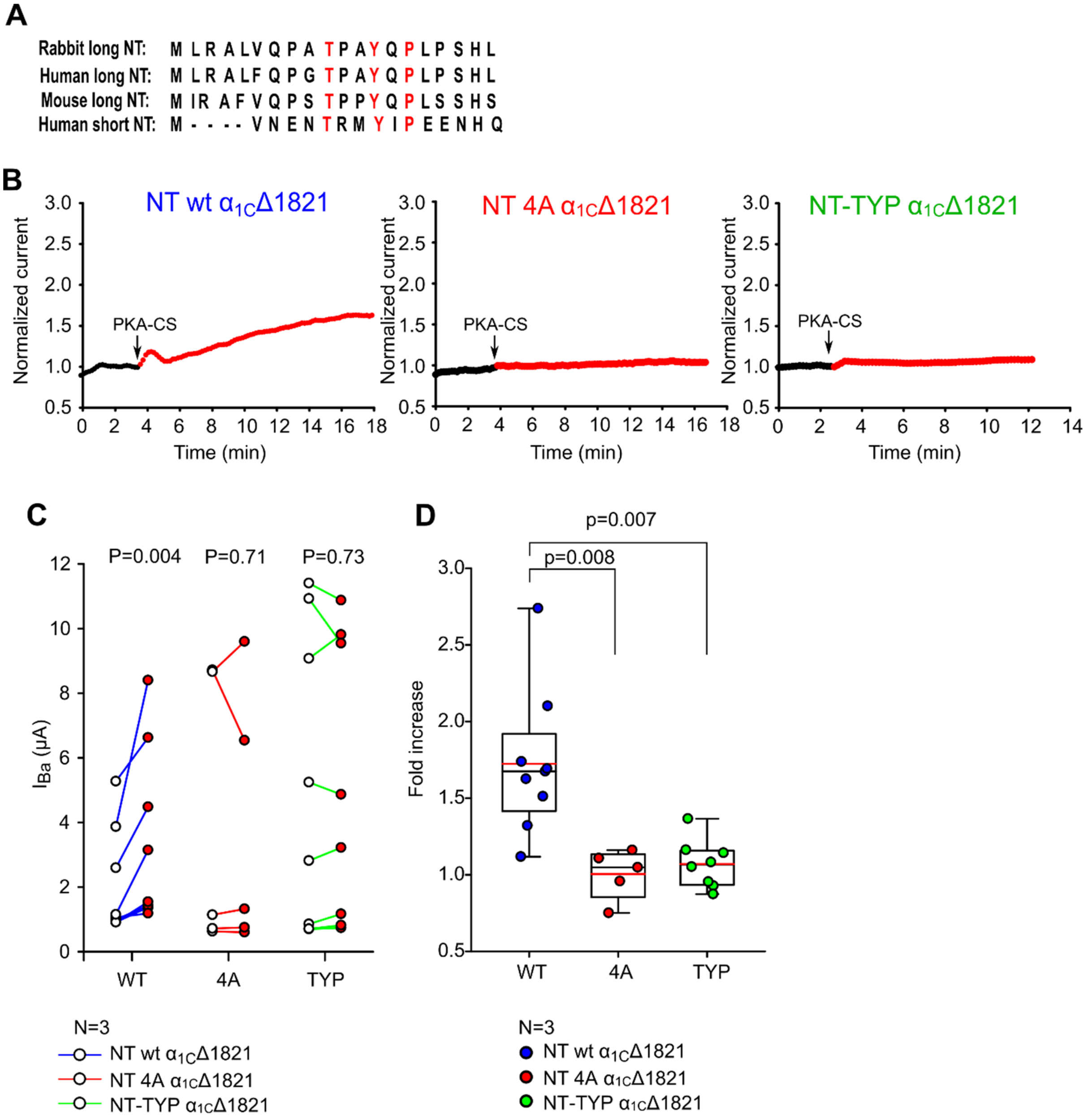


**Figure S1. Mutations within the initial segment of long-NT disrupt Rad-independent cAMP regulation of Ca$_V$1.2-α$_{1C}$Δ1821. A,** Amino acid sequences of the NH$_2$-terminal part of long-NT and short-NT isoforms of rabbit, human and mouse α$_{1C}$. Amino acids in positions T$_{10}$, Y$_{13}$ and P$_{15}$ (TYP motif, marked in red) are conserved in all isoforms. In this study only the long-NT isoforms were used. In these experiments we used mouse α$_{1C}$Δ1821[24] **B,** Representative diary plots of normalized Ba$^{2+}$ current at 20 mV. Figure show the time course of change in I$_{Ba}$ following the intracellular injection of 21 ng PKA catalytic subunit (PKA-CS, red trace). The channel was expressed in *Xenopus* oocytes in full subunit composition: α$_{1C}$Δ1821 (3 ng RNA), β$_{2b}$ (3 ng), α$_2$δ

(3 ng). In the left panel, the oocytes expressed mouse $\alpha_{1C}\Delta1821$ (NT wt α1CΔ1821). In the middle and right panels, mRNAs of mouse $\alpha_{1C}\Delta1821$ with alanine substitution of amino acids 2-5 (NT 4A $\alpha_{1C}\Delta1821$) and the TYP motif (NT-TYP $\alpha_{1C}\Delta1821$) were used, respectively. **C,** "before-after" plots of PKA-CS induced changes in $I_{Ba}$ in individual cells. N=3 experiments; statistics: paired t-test. **D,** Fold increase in $I_{Ba}$ induced by 21 ng PKA-CS. N=3; statistics: p=0.002, Kruskal-Wallis One Way ANOVA on Ranks followed by Dunnett's test.

*Fig. S2.*

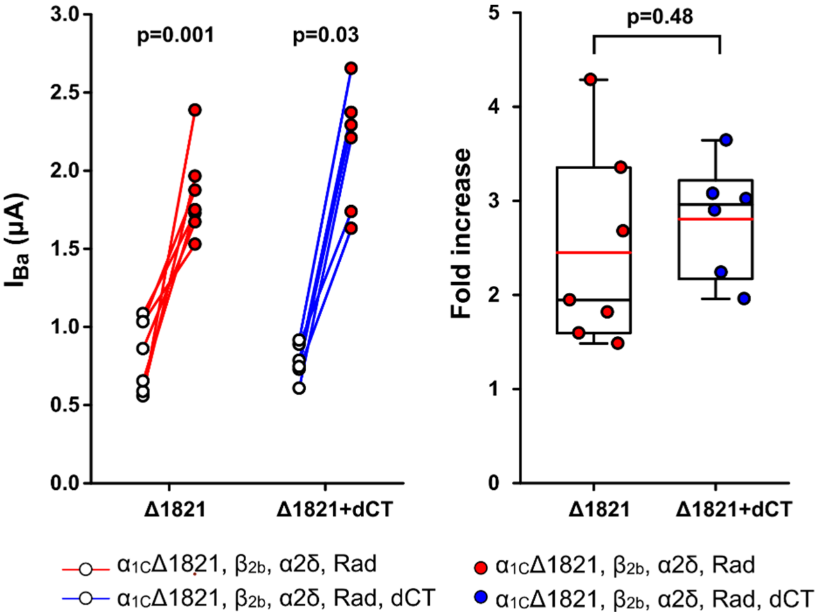


**Figure S2. The role of the clipped distal C terminus of $\alpha_{1C}$ in Rad-dependent regulation.** **A,** "before-after" plots of cAMP induced changes in $I_{Ba}$ in individual cells co-expressing $Ca_V$1.2-$\alpha_{1C}$Δ1821 and Rad, with or without the dCT (1822-2171) expressed as a separate protein[2424]. N=1 experiment; statistics: paired t-test (no dCT), Wilcoxon Signed Rank Test (with dCT). **B,** Fold increase in $I_{Ba}$ induced by cAMP injection. N=1; statistics: t-test.

***Fig. S3.***

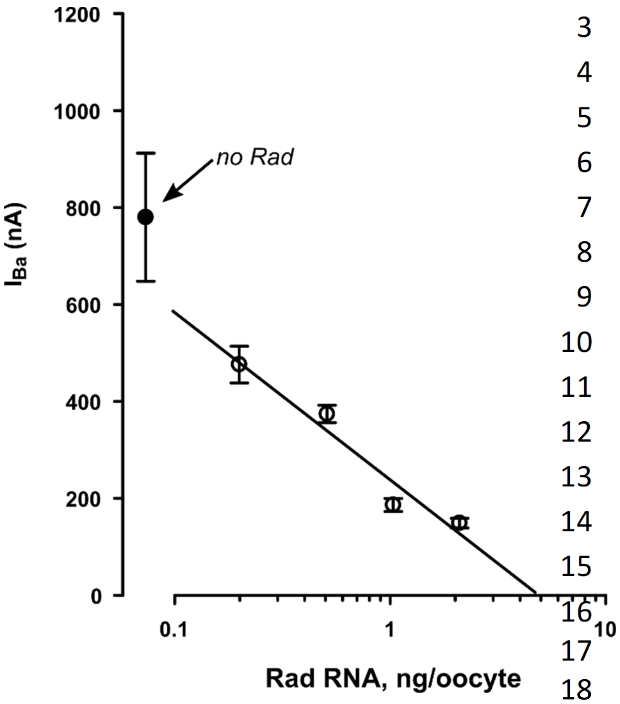


**Figure S3. Inverse correlation between Rad concentration and barium currents in full-length $\alpha_{1C}$.** Rad reduces the $Ba^{2+}$ current of $Ca_V1.2$-$\alpha_{1C}$ wt in a dose-dependent manner. The channel was expressed in *Xenopus* oocytes in full subunit composition: wt $\alpha_{1C}$ (5 ng RNA), $\beta_{2b}$ (5 ng), $\alpha_2\delta$ (5 ng). $I_{Ba}$ decreased with increasing doses of Rad RNA (Pearson correlation, r=-0.85, p=0.002). Each point presents mean±SEM from 4 to 10 oocytes (N=1 experiment). The linear regression line was drawn for non-zero doses of Rad.

#### Supplemental references (numbered as in main text)

14. Yang L, Katchman A, Samad T, Morrow J, Weinberg R and Marx SO. β-adrenergic regulation of the L-type $Ca^{2+}$ channel does not require phosphorylation of $\alpha_{1C}$ Ser1700. *Circ Res*. 2013;113:871-80. 10.1161/CIRCRESAHA.113.301926

24. Oz S, Pankonien I, Belkacemi A, Flockerzi V, Klussmann E, Haase H and Dascal N. Protein kinase A regulates C-terminally truncated $Ca_V1.2$ in *Xenopus* oocytes: roles of N- and C-termini of the $\alpha_{1C}$ subunit. *J Physiol*. 2017;595:3181-3202. 10.1113/JP274015

29. Hullin R, Singer-Lahat D, Freichel M, Biel M, Dascal N, Hofmann F and Flockerzi V. Calcium channel β subunit heterogeneity: functional expression of cloned cDNA from heart, aorta and brain. *EMBO J*. 1992;11:885-90.

30. Opatowsky Y, Chen CC, Campbell KP and Hirsch JA. Structural analysis of the voltage-dependent calcium channel β subunit functional core and its complex with the α1 interaction domain. *Neuron*. 2004;42:387-99. 10.1016/s0896-6273(04)00250-8
